# Bacterial persistence emerges mainly from antibiotic-specific networks rather than slow growth

**DOI:** 10.64898/2026.08.20.746038

**Authors:** François Beaufay, Camille Froment, Safia Zedek, Laurence Van Melderen

## Abstract

Bacterial persisters are rare phenotypic variants that survive antibiotic treatment and contribute to infection relapse and resistance emergence, yet the genetic basis of persistence remains only partially understood. Here, we developed a genome-wide Tn-seq approach to identify genes and pathways involved in persistence in exponentially growing *Escherichia coli* while avoiding persister enrichment and controlling for the confounding effects of growth. We found that persistence was largely independent of growth, and mutations that impaired growth could either increase or decrease survival. Furthermore, persistence was strongly antibiotic-specific. Ampicillin persistence relied primarily on pathways linked to energy metabolism, whereas ofloxacin persistence was associated on pathways involved in DNA repair and the maintenance of genome integrity. This study supports a model in which bacterial persistence is mainly an antibiotic-specific phenomenon governed by distinct physiological pathways rather than being universally associated with growth arrest.

## Introduction

The discovery of penicillin marked a major breakthrough in modern medicine, enabling the treatment of previously deadly bacterial infections [1]. Yet, alongside the rapid emergence of antibiotic resistance, early studies revealed that antibiotic treatment does not uniformly eradicate bacterial populations [2, 3]. In particular, penicillin failed to sterilize *Staphylococcus* cultures, uncovering a phenomenon known as bacterial persistence. Persister cells constitute rare subpopulations of genetically susceptible bacteria that transiently survive exposure to bactericidal antibiotics, thereby contributing to treatment failure and infection relapse [4, 5]. Unlike antibiotic resistance, persistence is reversible and non-heritable, emerging from phenotypic heterogeneity within clonal populations [6]. Persisters have classically been described as slow-growing or dormant cells that pre-exist within bacterial populations [7–9]. In this model, reduced metabolic activity, characterized by low ATP levels and limited biosynthetic activity, diminishes the activity of antibiotic targets and thereby promotes survival during treatment [7, 10–13]. Consistent with this view, persistence has been associated with reduced translation [14, 15], cytosol acidification [14, 16], and protein aggregation [17–19]. However, this strictly dormant model appears incomplete as persisters were also found to emerge from metabolically active and even dividing cells at the time of antibiotic exposure [20–23].

At the physiological level, persistence has been associated with diverse cellular pathways, including stress responses such as the RpoS-mediated general stress response and the stringent response, energy metabolism (notably the tricarboxylic acid (TCA) cycle, electron transport chain (ETC) and ATP synthesis) as well as proteostasis mediated by ATP-dependent proteases and chaperones and drug efflux [8, 24–26]. Despite these advances, comparisons across studies remain challenging because of substantial differences in experimental conditions, bacterial strains and persistence assays. Moreover, many studies rely on strategies that artificially increase the persister frequencies (10^−6^ to 10^−2^) in bacterial population including hyper-persistent or auxotroph mutants (e.g. MetG*, HipA7, adenine auxotroph) [15, 27, 28] or treatments that reduce growth and metabolism (*e.g*. stationary phase, nutrient limitation, pre-treatment with bacteriostatic antibiotics, serine hydroxamate or CCCP (carbonyl cyanide m-chlorophenylhydrazone), or overexpression of toxins or other regulatory proteins [14, 23, 29–31]. While these approaches have substantially advanced our understanding of persistence, the conditions used to increase the persister abundance may influenced the genetic landscape identified, with some determinants reflecting the specific experimental conditions rather than persistence *per se*. Similarly, previous genome-wide efforts to identify genes driving persistence have produced limited consensus [32–35]. The rarity and transient nature of persister cells, together with the difficulty of distinguishing them from growth-arrested or dead cells, have complicated their systematic characterization. As a result, key aspects of persister physiology remain unresolved, including whether persistence is associated with reduced or sustained metabolic activity and the extent to which the underlying mechanisms are shared across antibiotics.

Here, we applied a transposon sequencing (Tn-seq) based methodology to systematically identify genes associated with antibiotic persistence in exponentially growing *E. coli* wild-type cells, without relying on treatments or conditions that artificially increase persistence frequency. Additional steps of clearing and signal amplification not only enable the robust identification of genes promoting persistence but also genes causing severe persistence reduction close to the limits of detection. Finally, this approach allowed us to assess the relationship between growth and persistence for each non-essential gene of *E. coli*. By applying this framework to ampicillin and ofloxacin, two bactericidal antibiotics with distinct modes of action, we sought to determine whether persistence is governed by common cellular processes or by antibiotic-specific physiological pathways.

## Results

### Setting up a robust methodology to identify genes involved in persistence to antibiotics

A dense MG1655 *E. coli* transposon library comprising approximately 300,000 mutants was generated which corresponds to an average insertion every ∼15 bp across the genome. While high library diversity is paramount for Tn-Seq in general, the low frequency of persister cells introduces additional constraints and inherent limitations, including severe population bottlenecks (a random loss of genetic diversity due to drastic population reduction) and an increased risk of false-positive signals due to the low ratio of persister to dead cells. Therefore, we optimized key experimental parameters to establish a workflow enabling robust detection of persistence determinants to antibiotics in exponentially growing *E. coli* (Fig. 1A).

**Figure 1.**
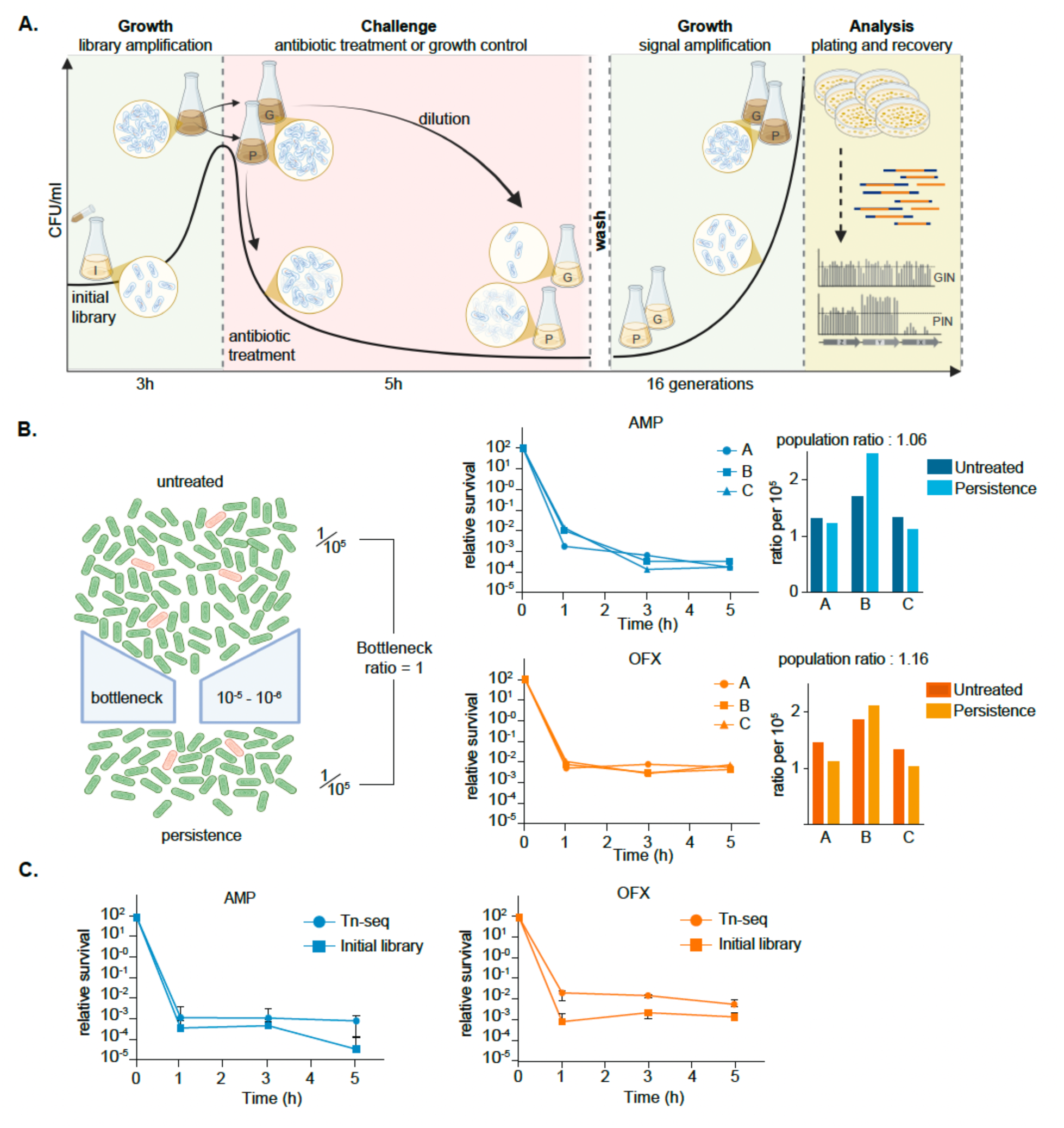
High-throughput Tn-seq approach for measuring growth and persistence indices in bacterial populations. (**A**) Experimental workflow. An *E. coli* transposon (Tn) library (I: initial library) was grown in LB medium to an OD_600nm_ of 0.8 (∼3 h) and exposed to antibiotic treatment for 5 h (P). Cells were then washed, resuspended in fresh medium, and allowed to regrow for ∼16 generations before plating on LB agar. Colonies were pooled, genomic DNA was extracted, and samples were subjected to sequencing. From the 0.8 OD_600nm_ initial library, a growth control (G) was processed in parallel, except that the antibiotic treatment was replaced by a dilution step (10⁻⁵-10⁻⁶) corresponding to the survival frequency observed after antibiotic exposure. Initial, growth control (G), and persister (P) libraries were sequenced. (**B**) Assessment of bottleneck effects. Mixed populations of red- and green-fluorescent cells (mScarlet/NeonGreen), mimicking the diversity of the Tn library, were exposed to ampicillin (AMP, upper panels) or ofloxacin (OFX, lower panels) for 5 h and plated on LB agar. Colonies recovered before treatment (0 h) and after treatment (5 h) were pooled, and the red-to-green cell ratio was quantified by flow cytometry. A bottleneck index close to 1 indicates preservation of population diversity and the absence of a detectable bottleneck effect (n = 3). (**C**) Time-kill curves of the initial Tn-Seq library (square) and persister-enriched fraction following the experimental workflow performed with AMP or OFX. Survival kinetics during treatment with AMP or OFX were determined for the initial Tn library (squares) and for the persister-enriched population (circles) used for DNA extraction and sequencing (n =3). (Blue = AMP; orange = OFX).

Because bacterial growth rate is a major determinant of antibiotic susceptibility [7, 36], we quantified persistence frequencies across different growth phases by starting antibiotic treatment at increasing OD_600nm_ (0.5, 0.8, 1, 1.5 and 2) (Extended Data Fig. 1A). Consistent with previous studies demonstrating that nutrient limitation and reduced growth promote survival to antibiotic [37, 38], we observed that persistence frequencies increased markedly as cultures approached stationary phase (OD_600nm_ > 1). This trend was observed following 5 h exposure to both ampicillin (AMP; 100 μg/mL, corresponding to 10-fold the MIC) and ofloxacin (OFX; 5 μg/mL, corresponding to 80-fold the MIC). We therefore selected an initial OD_600nm_ of ∼ 0.8, which provided optimal library representation while maintaining the characteristic biphasic killing curves and limiting the growth phase-dependent increase in persister frequency.

To assess potential population bottlenecks and random loss of Tn-mutants during the antibiotic treatment, we mixed cells constitutively expressing mScarlet or NeonGreen at ∼1:10⁵ ratio, approximating the Tn-library diversity. Following 5 h exposure to AMP or OFX, mixed populations were plated, collected and sorted based on fluorescence. Red-to-green ratios were comparable before and after antibiotic treatment, indicating the absence of detectable bottlenecks under these conditions (Fig. 1B, Extended Data Fig. 1B). Importantly, expression of the fluorescent proteins did not affect persistence to either antibiotic (Extended Data Fig. 1C). Then, to eliminate intact but dead cells that classically contaminate bulk/populational Tn-Seq, RNA-seq or proteome analyses, we incorporated a post-treatment outgrowth phase corresponding to approximately 16 generations, thereby reducing the contribution of residual DNA by an estimated 10^6^-fold (Fig. 1A). This step was particularly important for OFX-treated samples, as fluoroquinolone-mediated killing largely preserves cellular integrity which can delay the clearance of DNA from dead cells. Finally, to account for the impact of growth on mutant representation and distinguish persistence from general fitness effects, we included a growth-control condition to each experiment. In this control, antibiotic treatment was replaced by a dilution step matching the survival fraction observed after antibiotic exposure (∼ 10⁻⁵ – 10⁻⁶) (Fig. 1A). This approach allowed us to quantify growth-associated changes and, importantly, correct for their contribution to measure antibiotic specific effect in the persistence analysis.

Altogether, this workflow enabled the robust discrimination and quantification of growth fitness (initial library versus growth control) and persistence fitness (antibiotic-treated versus growth control) following exposure to AMP and OFX.

### Disruption of most of non-essential genes does not affect persistence

Applying the workflow described above (Fig. 1A), the Tn library was exposed to either AMP or OFX for 5 h, resulting in the characteristic biphasic killing kinetics associated with persistence. Upon regrowth, the surviving populations exhibited an approximately 10-fold increase in persister frequency when re-challenged with the corresponding antibiotic (AMP p=0.0021, OFX p < 0.0001), confirming that the initial treatment had indeed selected for genes associated with increased persistence (Fig. 1C). For each antibiotic, the workflow was performed in three independent biological replicates, yielding comparable levels of persistence, which were therefore combined for further analysis (Extended Data Fig. 1D). Transposon insertion sites were subsequently mapped across the *E. coli* MG1655 genome using custom bioinformatic pipelines [39] (Table S1). First, essential genes were identified based on the insertion index score, a methodology previously used to measure gene essentially in Tn-seq studies [40, 41]. Of the 4,494 genes encoded by MG1655, 553 and 572 were classified as essential in absence of antibiotic under the AMP and OFX experimental workflows, respectively (Table S2). The resulting essential gene sets were consistent with previously published genome-wide Tn-seq essentiality analyses [41] (Table S2), supporting the robustness of our dataset. We focused our analysis on the 3,941 (AMP) and 3,922 (OFX) non-essential genes of MG1655 (referred as genes hereafter). To quantify how each gene influence bacterial growth and persistence, we calculated two fitness indices. The growth fitness index (GIN) was defined as the log₂ ratio of the cumulative insertion frequency within a given gene in the growth-control population relative to the initial library, providing a measure of the fitness cost or benefit associated with disruption of that gene during growth (Table S3). The persistence fitness index (PIN) was defined as the log₂ ratio of the cumulative insertion frequency within a given gene in the antibiotic-treated population relative to the growth-control population, thereby eliminating the growth effect of each insertion and quantifying specifically the effect of gene disruption on persistence (Table S3). Positive GIN or PIN values indicate enrichment of insertion mutants under the corresponding condition, whereas negative values indicate depletion. Genes with GIN or PIN values greater than 1 or lower than -1 (GIN or PIN ∉]-1, 1[) were considered enriched or depleted, respectively. We considered smaller effect not biological meaningful and therefore discarded them from further analyses. Using this approach, both indices were determined for 3,941 and 3,922 genes under the AMP and OFX conditions, respectively, representing the majority of the ∼ 4,494 genes encoded by the MG1655 genome (Table S3). GIN analysis identified a relatively small subset of genes with substantial effects on growth, accounting for 6.8% and 8.0% of AMP and OFX gene datasets, respectively (Extended Data Fig. 2A). Although Tn mutants exhibiting reduced growth fitness (GIN ≤ -1) were similarly represented in both datasets (6.6% and 6.7% for AMP and OFX, respectively), Tn mutants with increased growth fitness (GIN ≥ 1) were more frequently identified in the OFX dataset (1.3%) than in the AMP dataset (0.22%) (Extended Data Fig. 2A). This discrepancy is likely attributable to experimental variability affecting a relatively small subset of mutants close to the biological threshold (Table S3). Nevertheless, GIN values remained strongly correlated between the two experiments (R_Pearson_ = 0.81, p < 0.0001), supporting the robustness and reproducibility of the growth index measurements (Extended Data Fig. 2B-C). PIN values identified a small fraction of genes with substantial effects on persistence, corresponding to 5.2% (208/3,941) and 4.0% (157/3,922) of genes under AMP and OFX conditions, respectively (Fig. 2A, B). Notably, AMP treatment predominantly identified mutants with reduced persistence (Fig. 2B), with 82% (170/208) of mutants exhibiting PIN values ≤ 1. In contrast, OFX treatment generated a more balanced distribution of phenotypes, with nearly equal proportions of mutants displaying increased persistence (PIN ≥ 1; 50.3%, 79/157) or decreased persistence (PIN ≤ -1; 49.7%, 78/157) (Fig. 2A-B). Interestingly, GIN values for mutants with altered PIN were highly correlated between OFX and AMP experiments (AMP: R_Pearson_ = 0.91, p < 0.0001; OFX: R_Pearson_ = 0.93, p < 0.0001,), supporting the interpretation that the observed differences in PIN are driven by persistence phenotypes rather than growth-related effects (Extended Data Fig. 2C). Altogether, these results indicate that only a relatively small subset of genes has a measurable impact on persistence and further suggest that the molecular basis underlying persistence are shaped by the mode of action of the antibiotic.

**Figure 2.**
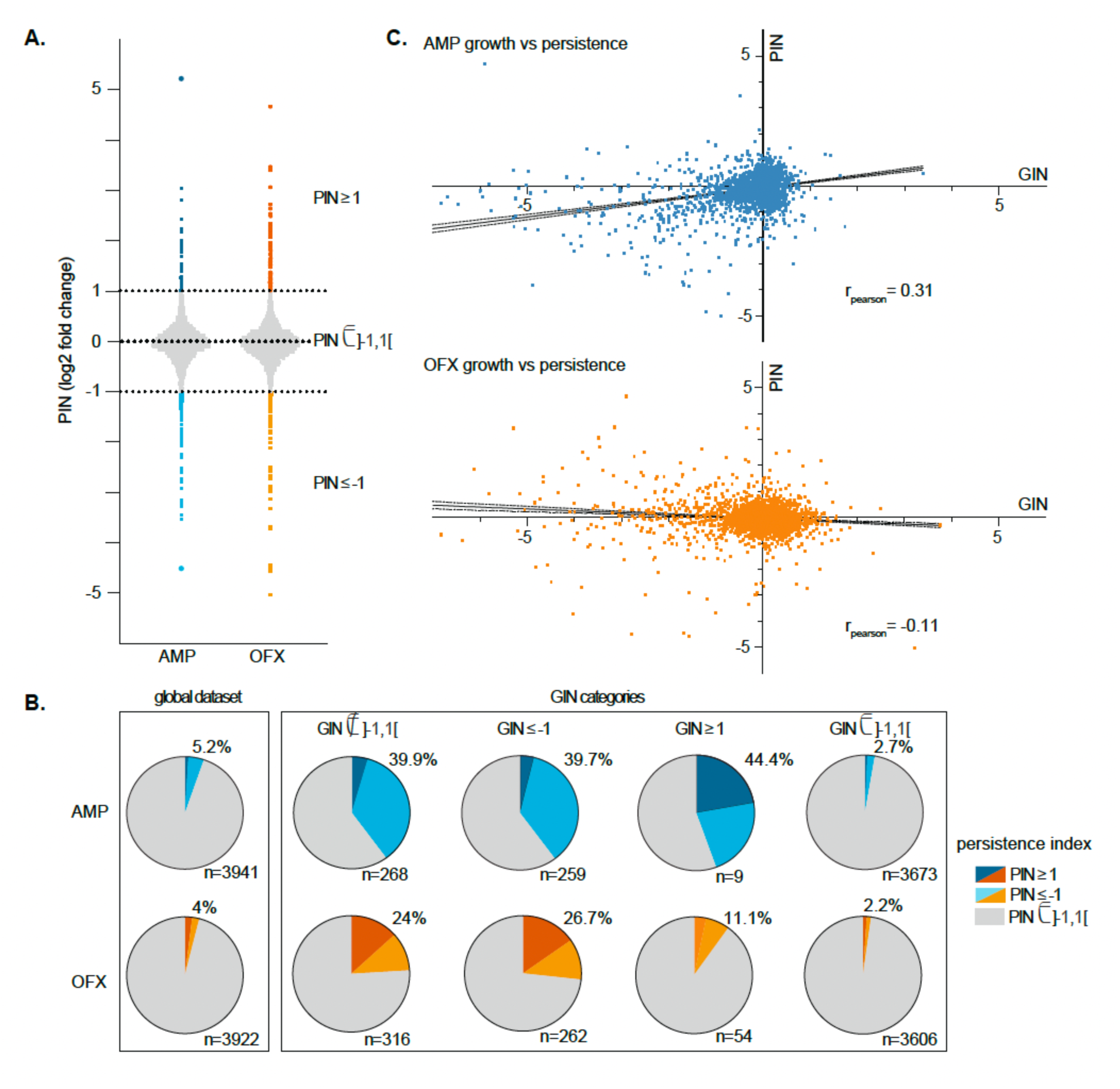
Genome-wide Tn-seq analysis of genes aIecting persistence. (**A**) Persistence index (PIN) of each non-essential gene upon OFX and AMP treatment. Significant threshold was set at +1 and -1. (**B**) Persister pie charts illustrating the link between growth (GIN) and PIN to AMP (upper panels) and OFX (lower panel). From left to right, distribution of PIN values for all non-essential genes in the global dataset (GIN ∉]-1, 1[), genes with positive and negative GIN values (GIN ∉]-1, 1[), with GIN negative values (GIN ≤ -1), with positive GIN values (GIN ≥ 1) and with non-significative GIN values (GIN ∈]-1, 1[). (**C**) Comparison of GIN and PIN values for each non-essential genes (AMP condition, upper panel; OFX condition, lower panel). Data are log_2_ fold change. Curve represents the fitted linear regression with the 95% confidence intervals. (Blue = AMP; orange = OFX).

### Perturbation of the growth rate generally affects persistence to both AMP and OFX

Persister cells have traditionally been viewed as metabolically inactive or dormant subpopulations characterized by reduced ATP levels and slow growth in comparison to the bulk population at the time of antibiotic exposure [6, 10, 12, 30]. Under this model, a strong relationship between GIN and PIN would be expected. Consistent with this hypothesis, persistence-associated genes were strongly overrepresented among mutants exhibiting altered growth fitness (GIN ∉]-1, 1[). Although only 5.2% and 4.0% of Tn mutants from the global dataset were significantly affected for persistence under AMP and OFX conditions, respectively, these proportions increased nearly eight- and six-fold, reaching 39.9% and 24.0% among mutants with significant growth alterations (GIN ∉]-1, 1[) (Fig. 2B). These results indicate that mutations affecting growth are more likely to influence persistence. However, despite this strong enrichment, correlation analysis revealed only weak associations between GIN and PIN (Fig. 2C). For the AMP condition, a modest positive correlation (R_Pearson_ = 0.315, R² = 0.099 p < 0.0001) indicates that growth explained only approximately 10% of the variation in persistence. Under OFX treatment, the association was even weaker (R_Pearson_ = - 0.111, R² = 0.012 p < 0.0001) with growth accounting for only approximately 1% of the variation in persistence. These data indicate that nearly 90% and 99% of the variation in persistence upon AMP and OFX treatment respectively is not explained by growth. Non-parametric permutation tests confirmed our observations (100,000 permutations for AMP and OFX conditions, p < 0.0001) supporting the conclusion that persistence is largely independent of bacterial growth under either antibiotic condition. Importantly, the impact of a gene disruption on growth did not predict whether it would promote or impair persistence. Under the OFX condition, approximately half of the mutants with reduced growth (GIN ≤ -1) exhibited enhanced persistence, whereas the other half displayed reduced persistence. This proportion remained essentially unchanged compared to the global dataset (for the GIN ≤ -1 dataset, n=262, 42 genes with PIN ≥ 1 and 34 with PIN ≤ -1; for the global dataset, n=3922, 79 genes with PIN ≥ 1, and 78 with PIN ≤ -1) (Fig 2B). It was even more striking under the AMP condition, where the majority of the mutants with reduced growth (GIN ≤ −1) exhibited reduced persistence, ruling out any correlation between slow growth and enhanced persistence (for the GIN ≤ -1 dataset, n= 259, 38 genes with PIN ≥ 1 and 170 with PIN ≤ -1; for the global dataset, n=3941, 12 genes with PIN ≥ 1 and 95 with PIN ≤ - 1). Persistence phenotypes were also observed among mutants exhibiting increased growth (GIN ≥ 1) (Fig. 2B-C). Under OFX condition, as observed for growth-impaired mutants, these genes were nearly equally divided between enhanced and reduced persistence phenotypes (for the GIN ≥1 dataset, n=54, 2 genes with PIN ≥ 1 and 4 with PIN ≤ -1), further indicating that the persistence phenotype is independent of growth. The number of mutants with increased growth identified in the AMP dataset (n = 9) was too small to allow meaningful interpretation. Finally, a subset of mutants displayed significant persistence phenotypes despite having no detectable effect on growth (GIN ∈]-1, 1[), accounting for 2.7% and 2.2% of genes in the AMP and OFX datasets, respectively (Fig. 2B). These findings demonstrate that persistence can be genetically uncoupled from growth.

Collectively, these results challenge the prevailing view that slow growth *per se* promotes persistence. Instead, our finding suggest that growth perturbations increase the likelihood of altering persistence, with the direction and magnitude of the persistence phenotype being determined by the specific cellular functions affected rather than by growth itself.

### Individual deletion mutant analysis validates the high throughput workflow

To validate the Tn-Seq data, individual deletion mutants for a subset of candidate genes were constructed (Fig. 3A). Growth, MIC, and persistence to AMP and OFX were assessed individually (Fig. 3A, Extended data Fig. 3). Even though Tn-seq-derived GIN values reflect a composite fitness measure integrating lag phase, growth rate, and competition for resources, whereas the growth of individual deletion mutants was quantified independently by OD_600nm_ measurements (Extended Data Fig. 3A-B), both approaches produced comparable estimates of growth fitness. Indeed, GIN values derived from Tn-seq correlated well with those calculated from the deletion mutants (R_Pearson_ = 0.71, R² = 0.5 p < 0.0001 and R_Pearson_ = 0.74, R² = 0.57 p < 0.0001for AMP and OFX, respectively) Extended Data Fig. 3C). In addition, GIN values derived from the Tn-seq datasets under AMP and OFX conditions were highly correlated (R_Pearson_ = 0.92, R² = 0.85, p < 0.0001), demonstrating the high reproducibility of the GIN measurements across experimental conditions.

**Figure 3.**
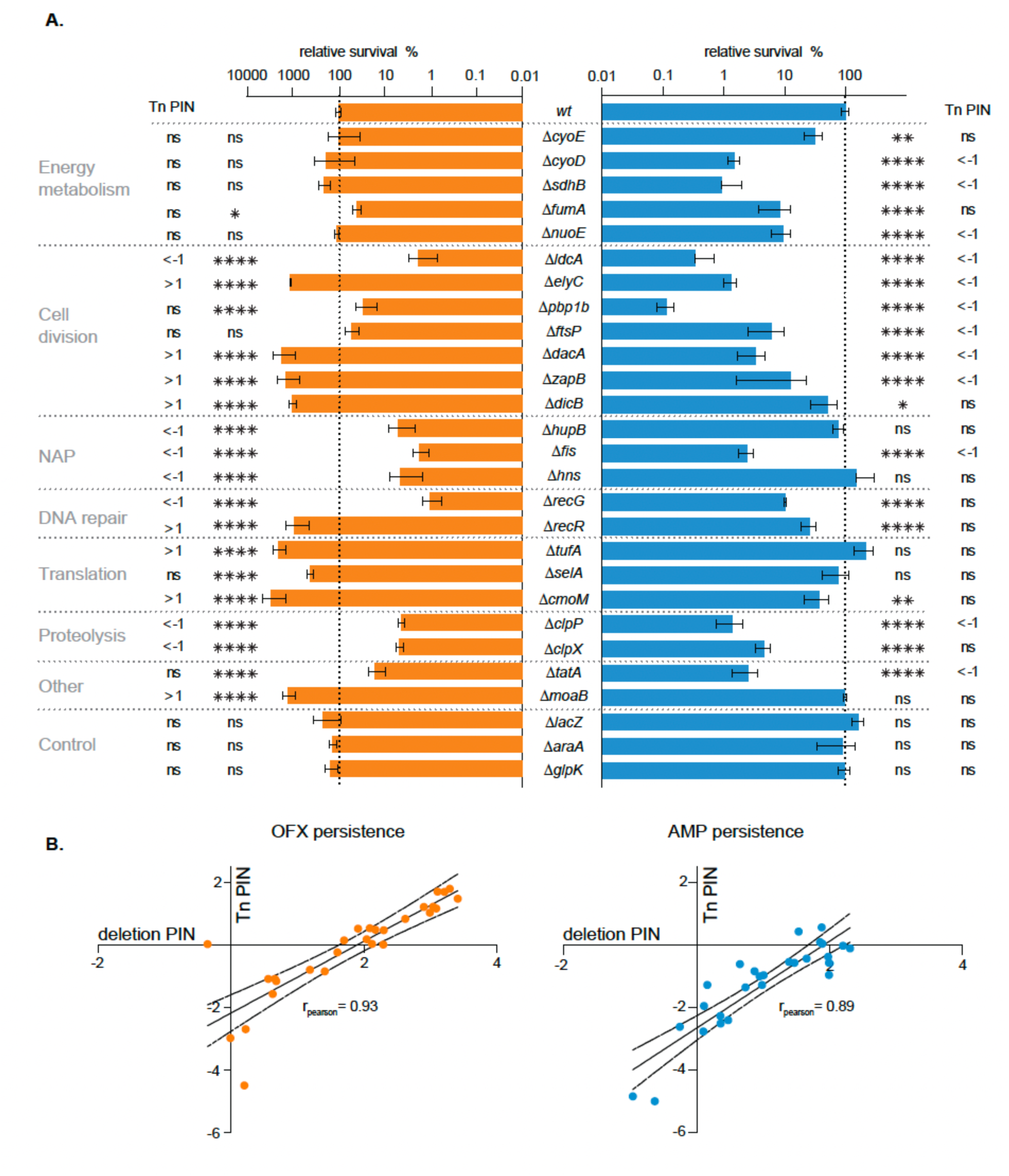
Persister assays of selected gene deletion mutants confirmed the high-throughput approach. (**A**) Survival of deletion mutants relative to the WT strain after 5 h treatment with OFX (orange) or AMP (blue). WT survival was set at 100%. Genes are organized according to their functional annotations. For each gene, experiments were performed in triplicates (for biological replicates and killing curve, see Fig. S3) (n ≥ 3; * p<0.05, ** p<0.01, *** p<0.001, **** p<0.0001, ns = non-significative; one-way ANOVA). For each deletion mutant, the PIN category of the corresponding Tn-mutant is indicated. (**B**) Comparison of PIN values between the Tn-mutants and their corresponding deletion mutants (OFX, left panel; AMP, right panel). Data are expressed in log_2_ fold change. Curve represents the fitted linear regression with the 95% confidence intervals.

More importantly, all tested deletion mutants recapitulated the persistence phenotypes observed in the Tn-Seq analysis (Fig. 3A, Extended Data Fig. 3D). PIN values derived from deletion mutants were highly correlated with those obtained by Tn-seq, although not always reaching statistical significance individually (for AMP R_Pearson_ = 0.89, R^2^ = 0.79, p < 0.0001; for OFX R_Pearson_ = 0.93, R^2^ = 0.86, p < 0.0001) (Fig. 3B). Moreover, except for *recR* and *recG* deletions displaying a 10-fold higher sensitivity to OFX, all mutants displayed MICs similar to the wild-type, confirming that the identified genes modulate persistence rather than resistance (Extended Data Fig. 3E). Furthermore, survival of the mutants in the presence of the antibiotics display the classical biphasic curves, supporting that the genes are involved in persistence (Extended Data Fig. 3F).

These findings demonstrate the robustness and accuracy of our Tn-seq approach for uncovering antibiotic-specific determinants of bacterial persistence.

### Distinct cellular pathways govern persistence to different antibiotics

To identify specific pathways associated with persistence, Tn-mutants with significantly altered PIN values (PIN ∉]-1, 1[) were grouped according to their functional annotation [42, 43] (Fig. 4, Table S4). Under AMP treatment, oxidative phosphorylation and cell division showed the strongest enrichment, followed by carbon metabolism, response to antibiotics, DNA repair, secretion systems, cell wall biogenesis, translation, and transcription regulation (Fig. 4A, Extended data Fig. 4A, Table S4). Protein network analysis using STRING was performed to explore how the different pathways associated with persistence are interconnected [44]. The resulting network revealed major modules centered on energy metabolism, encompassing oxidative phosphorylation and carbon metabolism; cell division/cell envelope biogenesis; and global regulation, comprising transcription factors, nucleoid-associated proteins (NAPs), and proteases (Extended Data Fig. 4B-C, Table S4). The energy metabolism module further encompassed proteins involved in iron homeostasis (Fur) and iron-sulfur cluster biogenesis (NfuA and IscA), consistent with the role of iron-sulfur cofactors in oxidative phosphorylation. Additional, smaller modules were associated with chromosome integrity, translation, response to antibiotics and secretion systems (Extended data Fig. 4B). Notably, gene disruption across these modules almost uniformly resulted in reduced persistence (PIN ≤ −1), with only few exceptions (Extended Data Fig. 4C, Table S4). No correlation was observed between GIN and PIN values in the different modules, further supporting the conclusion that persistence is uncoupled from growth-related fitness (Extended Data Fig. 4D).

**Figure 4.**
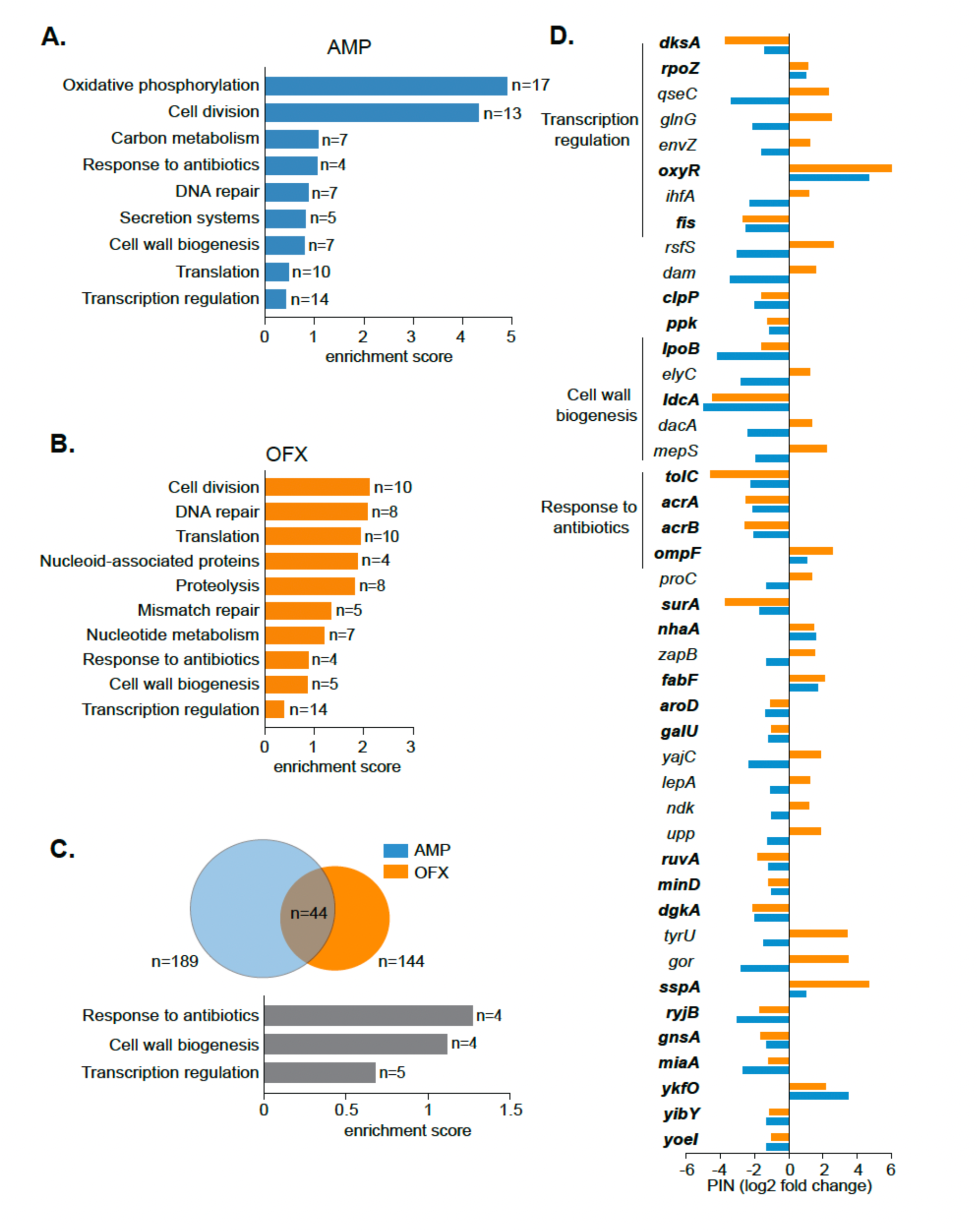
Main pathways aIecting bacterial persistence to ampicillin and ofloxacin. (**A-B**) DAVID functional classification of genes involved in persistence to AMP (**A**) and OFX (**B**). n represents the number of gene in each category. (**C-D**) Shared genes and pathways associated with persistence to AMP and OFX. (**C**) Venn diagram showing genes associated with persistence to AMP (blue) and OFX (orange). DAVID functional classification of the 44 genes associated with persistence to both antibiotics. (**D**) PIN values for AMP (blue) and OFX (orange) for the 44 shared genes. Genes are grouped according to their functional annotations, with bold indicating genes displaying the same PIN category under both antibiotic treatments. Data represent log_2_ fold changes derived from the Tn-seq analysis.

In contrast to AMP, OFX-associated genes were distributed across a broader number of functional categories with comparable enrichment scores including cell division, DNA repair, translation, NAPs, proteolysis, mismatch repair, nucleotide metabolism, response to antibiotics, cell wall biogenesis and transcription regulation (Fig. 4B, Extended Data Fig. 5A, Table S4). STRING analysis organized proteins from theses enriched pathways into interconnected cellular modules associated with chromosome integrity (DNA repair, NAPs, mismatch repair and nucleotide metabolism), cell division/cell envelope biogenesis (cell division, cell wall biogenesis, proteolysis) and global regulation (NAPs, transcriptional regulation and proteolysis) (Extended Data Fig. 5B-C, Table S4). Disruption of genes associated to these modules had divergent effects on persistence, even within the same functional category, with some mutations increasing persistence (PIN ≥ 1) and others decreasing it (PIN ≤ −1) (Extended Data Fig. 5C). As for AMP, we did not observe any correlation between PIN and GIN values when focusing on each module individually (Extended Data Fig. 5D).

Although several functional categories were significantly enriched in both AMP and OFX datasets, the specific genes within these shared categories were mostly different between antibiotics with only 44 genes common to both antibiotic (Fig. 4C; Extended Data Fig. 4C, 5C (common genes indicated in bold); Table S5). Gene enrichment analysis highlighted 3 functional categories namely response to antibiotics, cell wall biogenesis and transcription regulation (Fig. 4C, Extended Data Fig. 6A). Among these, approximately 60% (27/44) displayed consistent persistence phenotypes across both treatments, including 19 mutants with reduced persistence and 8 with increased persistence, albeit often with different magnitudes of effect (Fig. 4D, indicated in bold). STRING analysis revealed 3 main modules centered on response to antibiotics, cell division/cell envelope biogenesis and global regulation (Extended Data Fig. 6B-C). The only functional category showing a consistent increase in persistence across both antibiotic treatments was the response to antibiotics (R_Pearson_ = 0.96, R^2^ = 0.92, p = 0.04) (Extended Data Fig. 6C-D), represented by genes encoding the AcrAB-TolC multidrug efflux system and the outer membrane porin OmpF know to mediate the entry of various antibiotics including β-lactams and fluoroquinolones [45, 46] (Fig. 4D). In contrast, PIN values under AMP and OFX showed no correlation for the cell division/cell envelope biogenesis category (R_Pearson_ = 0.17). Within the global regulation category, several genes contributed to persistence under both antibiotic conditions, including genes involved in proteolysis, transcriptional regulation, and nucleoid organization (R_Pearson_ = 0.69) (Extended Data Fig. 6C-D). Note that GIN values were strongly correlated between the two antibiotic conditions (R_Pearson_ = 0.96). These findings indicate that, while a limited set of core cellular functions contributes to persistence across conditions, the genetic architecture of persistence is primarily dictated by the antibiotic’s mode of action rather than by a universal persistence mechanism.

## Discussion

Despite being recognized for more than eight decades [2, 3], the genetic basis of bacterial persistence remains incompletely understood. Progress in the field has been hampered by the intrinsic properties of persister cells. Indeed, persister cells are rare, transient phenotypic variants that arise within genetically identical populations and survive antibiotic treatment without acquiring heritable resistance mutations [4, 47]. To address these challenges, we developed a Tn-seq approach specifically designed to identify genes and pathways associated with persistence in exponentially growing populations. Importantly, our strategy avoids the genetic or environmental perturbations commonly used to enrich persister cells. Such perturbations may obscure the pathways contributing to persistence under unperturbed growth conditions or preferentially reveal pathways associated with triggered persistence, such as those operating during stationary phase [4, 6].

One of the central conclusions of this work is that persistence cannot be explained by reduced growth or metabolism alone. This observation was consistent for AMP, a β-lactam which is largely ineffective against non-dividing cells [36], and OFX, a fluoroquinolone whose activity is reported to be less growth-dependent than ampicillin [48]. Although growth-phenotypes were enriched among persistence associated genes, no correlation was observed between growth (GIN) and persistence (PIN) (Fig. 2C). Mutations causing comparable growth defects often had opposite effects on persistence, and several persistence-associated mutants exhibited no measurable growth defect. Together, these findings indicate that reduced growth is neither necessary nor sufficient for persistence. Instead, they indicate that the establishment or maintenance of the persister state emerges from molecular pathways that are largely independent of growth rate itself. Furthermore, our findings argue against the existence of a universal, common mechanism for persistence and instead point to antibiotic-specific pathways that are closely related to the mode of action of each antibiotic. Consistent with previous studies, OFX persistence was strongly associated with DNA repair functions [20, 31, 49, 50]. Importantly, our analysis extended this view by revealing a broader network of pathways involved in the maintenance of genome integrity, encompassing DNA repair, mismatch repair, nucleoid-associated proteins (NAPs), and nucleotide metabolism (Fig. 4, Extended Data Fig. 5). Given that OFX induces DNA double-strand breaks by poisoning DNA-gyrase complexes, the contribution of pathways involved in DNA repair, genome organization, and nucleotide homeostasis is consistent with the cellular damage caused by the antibiotic. In contrast, AMP persistence was strongly linked to energy metabolism particularly central carbon metabolism and oxidative phosphorylation (Fig. 4, Extended Data Fig. 4). The enrichment of genes involved in oxidative phosphorylation and ATP production indicates that survival to β-lactam requires the maintenance of sufficient energetic capacity [51]. Thus, rather than being associated with metabolic dormancy, persistence to AMP appears to rely on specific metabolic functions that support cellular maintenance during antibiotic exposure and/or subsequent recovery. Since PIN values derived from our Tn-seq screen represent a composite measure integrating the ability to withstand antibiotic exposure, exit from the persister state, and post-treatment recovery, the genes and pathways identified here may act at one or more of these distinct stages. For OFX, temporal information is available for the SOS response, which reaches maximal induction during recovery and appears to be primarily required after antibiotic removal [20, 52]. This suggests that at least part of the contribution of DNA repair functions identified in our screen may occur during post-treatment recovery. However, other OFX-associated pathways may act at different stages; for instance, nucleotide metabolism could contribute during antibiotic exposure by maintaining nucleotide pools required to cope with OFX-induced DNA damage. Similarly, the temporal contribution of the pathways associated with AMP persistence remains to be established. Dissecting when these antibiotic-specific pathways act will therefore be important to understand how their combined effects ultimately determine the final survival phenotype. Although our data highlight the strong antibiotic specificity of persistence pathways, several cellular processes were shared between AMP and OFX. Notably, the ‘response to antibiotics’ pathway including the outer membrane porin OmpF and the AcrAB-TolC efflux pump, showed similar effects on persistence across both treatments (Fig. 4, Extended Data Fig. 6). Consistent with previous studies, deletion of the *acrAB* genes decreased persistence to both OFX and AMP, whereas deletion of *ompF* increased persistence [24, 53, 54]. These common effects are consistent with the respective roles of AcrAB-TolC and OmpF in controlling intracellular antibiotic accumulation, a process expected to influence survival independently of the antibiotic mode of action. Moreover, increased AcrAB activity has been associated with enhanced survival during prolonged antibiotic exposure and increased mutagenesis, potentially facilitating the subsequent emergence of antibiotic resistance [55]. Beyond these 4 genes, we identified several cellular pathways including cell division, cell envelope biogenesis and translation that acts as recurrent hubs within persistence networks. However, their effects were not necessarily conserved between antibiotics. Similarly, global cellular regulators, comprising transcription factors, NAPs and proteases, could either promote or impair persistence depending on the antibiotic considered. These regulators may therefore act as connecting hubs between otherwise distinct cellular pathways, integrating antibiotic-specific physiological responses into broader persistence networks. Collectively, our findings support a model in which persistence primarily relies on distinct, antibiotic-specific physiological networks rather than on a universal survival mechanism associated with slow growth, reduced metabolic activity or dormancy. Beyond defining persistence networks for AMP and OFX, our study provides a general framework for systematically dissecting the genetic and physiological basis of persistence across antibiotics, paving the way toward a mechanistic understanding of antibiotic-specific survival strategies.

## Methods

### Bacterial strains and growth conditions

Strains used in experiments are all derivatives of the *E. coli* MG1655 laboratory strain ([56]. Strains, plasmids and oligonucleotides used in this study are listed in Table S6. Gene deletions were generated by λ red-mediated site-specific recombination [57] and confirmed by PCR. Chromosomal gene loci were transferred by phage P1 transduction to generate the final strains. Antibiotic cassettes were removed using site-specific recombination induced by expression of the FLP recombinase from the pCP20 plasmid. WT and mutant *E. coli* strains were grown at 37°C in Luria Bertani (LB) medium [tryptone (10 g/L), yeast extract (5 g/L), and NaCl (10 g/L) - Thermo Fisher Scientific]. The following antibiotics were added when appropriate: chloramphenicol (30 µg/mL), kanamycin (50 µg/mL), ampicillin (100 µg/mL) or ofloxacin (5 µg/mL).

### Persistence assays

Persistence assays were performed as described previously [58]. Briefly, overnight cultures grown in LB were diluted in fresh LB medium. Cells were treated with either OFX (5 μg/mL) or AMP (100 μg/mL) at an OD_600nm_ of 0.5. Samples were collected at 0, 0.5, 1, 3 and 5 h, washed, serially diluted in LB and plated on LB agar plates. Survival frequency was calculated as the ratio of the number of colony-forming units (CFUs) at a given time to the number of CFUs before antibiotic treatment. Survival of each mutant was normalized to the corresponding WT survival and expressed as a percentage. Experiments were performed in biological triplicates.

### Minimal inhibitory concentration

The MIC of OFX and AMP was determined using the agar dilution method as described [59]. Briefly, cultures were grown in LB and spotted on LB agar plates containing increasing concentration of AMP or OFX. MIC for each mutant was set at the lowest concentration preventing bacterial growth. Experiments were performed in biological triplicates.

### Growth curve

Overnight cultures grown in LB were diluted in 2mL of fresh medium at OD_600nm_ ∼ 0.05 and placed in clear bottom 24-well plates. Plates were inserted into an automated microplate reader and the OD_600nm_ was measured every 5 min for 24 h while being shaken at 140 rpm at 37°C. Lag time was measured as the time necessary for each mutant to reach 0.8 OD_600nm_ and exponential growth rate was extracted from the fitting curve of the ln(OD_600nm_). Values were normalized against the WT internal control for each experiment with WT value set at 100%. Experiments were performed in biological triplicates.

### Bottleneck assessment

MG1655 cells containing the plasmid-encoded mScarlet or mNeonGreen reporters were mixed at an approximate ratio of 1:10^5^ (red/green) and fluorescence was measured by flow cytometry before and after persistence assays with AMP and OFX at 100 and 5 μg/mL, respectively. Samples were diluted in PBS to an OD_600nm_ ∼ 0.01 and processed by an Attune NXT flow cytometer (Thermo Fisher Scientific) at a flow rate of 12.5 μL/min. In each experiment, a total of 10,000,000 events was analyzed per replicate. For each replicate, the ratio of red/green cells before and after antibiotic treatments was calculated. Comparison of both values was defined as the bottleneck score. A bottleneck score close to 1 indicates that cells initially present at a frequency of ∼10⁻⁵ remained proportionally represented following antibiotic treatment, arguing against a treatment-induced bottleneck capable of substantially distorting mutant representation.

### Tn-Seq methodology and analysis

A mini-Tn5 (EZ-Tn5™ <KAN-2>Tnp Transposome™ Kit - Lucigen) was introduced in the MG1655 strain by electroporation and plated on LB agar supplemented with kanamycin. At least 300,000 colonies were collected in LB with 10% glycerol and stored at -80°C to constitute the initial library. The initial library was diluted in LB medium and grown for ∼ 3h up to an OD_600nm_ ∼ 0.8. The culture was divided into a growth (G) and a persistence (P) fraction. The P fraction was treated with antibiotics for 5 h, centrifuged and cells were resuspended in LB fresh medium. The G fraction was not treated with antibiotics and was directly centrifuged and diluted 10^6^-fold (AMP) or 10^5^-fold (OFX) into LB fresh medium, corresponding to the survival rate of P cells under the corresponding antibiotic treatment. G and P cells were grown for 16 generations and plated onto large LB agar plates supplemented with kanamycin. Colonies were collected in LB with 10% glycerol and stored at -80°C. Genomic DNA from the initial library, P and G conditions for both antibiotics was extracted following manufacturer protocol (N-Qiagen) and resuspended in 50 µL elution buffer (5 mM Tris-HCl [pH 8.5]). Seventy-five base single end sequencing was performed on an Illumina NovaSeq (Fasteris, Switzerland). Reads corresponding to the mini-Tn5 insertion sites were first filtered with the 5′ -ACCTACAACAAAGCTCTCATCAACC Tn5 sequence before being processed as described in [39]. Over 150 million filtered reads for each P condition (AMP and OFX) were mapped onto the reference genome of the *E. coli* MG1655 strain (NC_000913) and converted to Sequence Alignment/Map (SAM) format using the Burrows-Wheeler Aligner and SAMtools, respectively, from the Sourceforge server (https://sourceforge.net/). Next, the number of reads overlapping each genomic position was computed using custom Python scripts. The total number of reads for internal 80 % of each open reading frame (ORF) (ES-read) in each condition was then computed using custom Python scripts. For comparison, the total number of reads was normalized to 1 million reads for each condition (ES-RPM) (Table S1). To extract Growth index values (GIN), for each gene, ES-RPM from growth libraries to each antibiotic was normalized to the ES-RPM of the initial library. To extract Persistence index (PIN), for each gene, ES-RPM from persistence libraries to each antibiotic was normalized to the ES-RPM of the growth library dissociating growth and persistence. Genes with a log_2_ fold change ≥ 1 or ≤ -1 were considered as genes of interest.

### Essential gene prediction

We rely on the insertion index score to determine if a gene was essential or non-essential as previously described [41]. Briefly, the insertion index score was obtained by normalizing the numbers of unique insertions in a gene by the size of the gene (in bases). To determine essentiality, we focus on unique insertions from growth control sequencing performed for AMP and OFX experiments. For each growth library, the frequency of insertion index score for each gene was plotted in a histogram using the Freedman-Diaconis rule for choice of bin widths (see Table S2). The frequency distribution of the insertion index scores was bimodal with genes to the left (low number of Tn insertions - exponential distribution) representing essential or severe fitness cost genes and genes to the right regrouping non-essential gene (high number of Tn-insertions - gamma distribution). Intersection between both distributions defined the essential threshold. For both workflow, genes with an insertion index score in the growth dataset ≤ 0.0488 (AMP,) and ≤ 0.0476 (OFX) were considered as essential. In addition, a small number of non-essential genes in our growth libraries displayed an extremely low number of unique insertions in our initial in which they were flagged as essential (10 genes for AMP; 5 for OFX). These genes were also classified as essential and removed from further analysis.

### Gene enrichment analysis

Functional annotation clustering was performed using the Database for Annotation, Visualization and Integrated Discovery (DAVID) against various database [42, 43]. Genes with 1 < PIN < -1 was compared against all non-essential genes of each Tn-library. DAVID uses a modified one-tailed Fisher’s Exact test (EASE), with a p-value < 0.05 considered significant to evaluate whether the number of genes associated with a given annotation term is greater than expected by chance. Then, DAVID groups related annotation terms together and assigns an Enrichment Score (the geometric mean (in -log scale) of member’s p-values in a corresponding annotation cluster) to the entire cluster.

Protein-protein interaction network analysis was performed using the STRING database (Search Tool for the Retrieval of Interacting Genes/Proteins, [44]. Genes with PIN > 1 and < -1 were mapped to corresponding proteins and interaction networks were constructed based on experimentally validated interactions, curated pathway knowledge, co-expression, genomic context, and computational predictions. Proteins were represented as nodes and interactions as edges. The combined score was computed by combining the probabilities from the different evidence channels and corrected for the probability of randomly observing an interaction. Network clustering was performed using the Markov Cluster Algorithm (MCL) implemented in STRING to identify groups of highly interconnected proteins that may participate in related biological functions.

### Statistical analyses

All the statistical analyses were performed using GraphPad Prism 9 software. A p-value < 0.05 was considered as significant.

## Acknowledgments

We are grateful to Kenny Petit for helping with the Tn-Seq data analysis. We thank Abel Garcia-Pino, Frédéric Goormaghtigh, Dukas Jurenas and Thierry Oms for careful reading and comments on the manuscript.

## Author information

These authors jointly supervised this work: François Beaufay, Laurence Van Melderen

### Contributions

L.V.M. and F.B. conceived and designed the experiments. F.B., C.M. and S.Z. performed the experiments. F.B. C.F. and L.V.M. analyzed data. L.V.M. and F.B. wrote the paper. The figures were prepared by F.B. All authors edited and revised the manuscript.

### Ethics declaration

The authors declare no competing interests

---

**Extended Data Figure 1.**
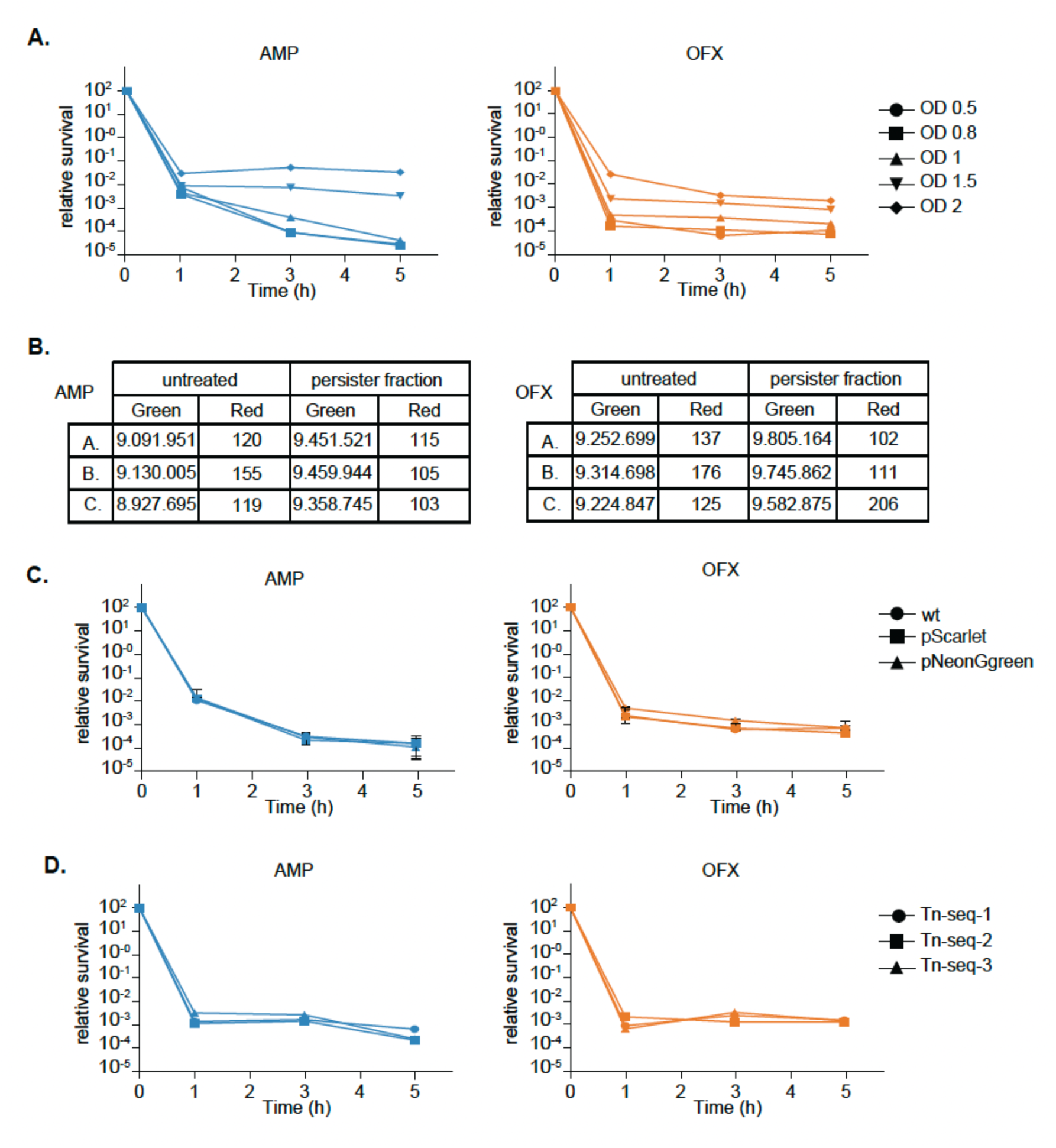
Detailed experimental set up to avoid a bottleneck eIect. (**A**) Optimization of the initial cell density before antibiotic treatment. Time-kill curves under AMP or OFX treatment with cultures at different OD_600nm_. (**B-C**) Assessment of bottleneck effects. The Tn-seq workflow was mimicked using cells constitutively expressing either a red or a green fluorescent protein. Red and green cells were mixed at an approximate ratio of 1:10⁵, corresponding to the diversity of the Tn library, and subjected to the AMP or OFX persistence assay described in Fig. 1B. The red-to-green ratio was quantified by flow cytometry before and after antibiotic treatment from three independent experiments (n = 10⁶ cells analyzed cells per sample). (**C**) No effect of mScarlet or NeonGreen fluorescent markers on AMP or OFX persistence frequency. Time-kill curves are shown for three independent experiments. (**D**) Survival kinetics during treatment with AMP or OFX from individual biological replicate used for DNA extraction and sequencing. (Blue = AMP; orange = OFX).

**Extended Data Figure 2.**
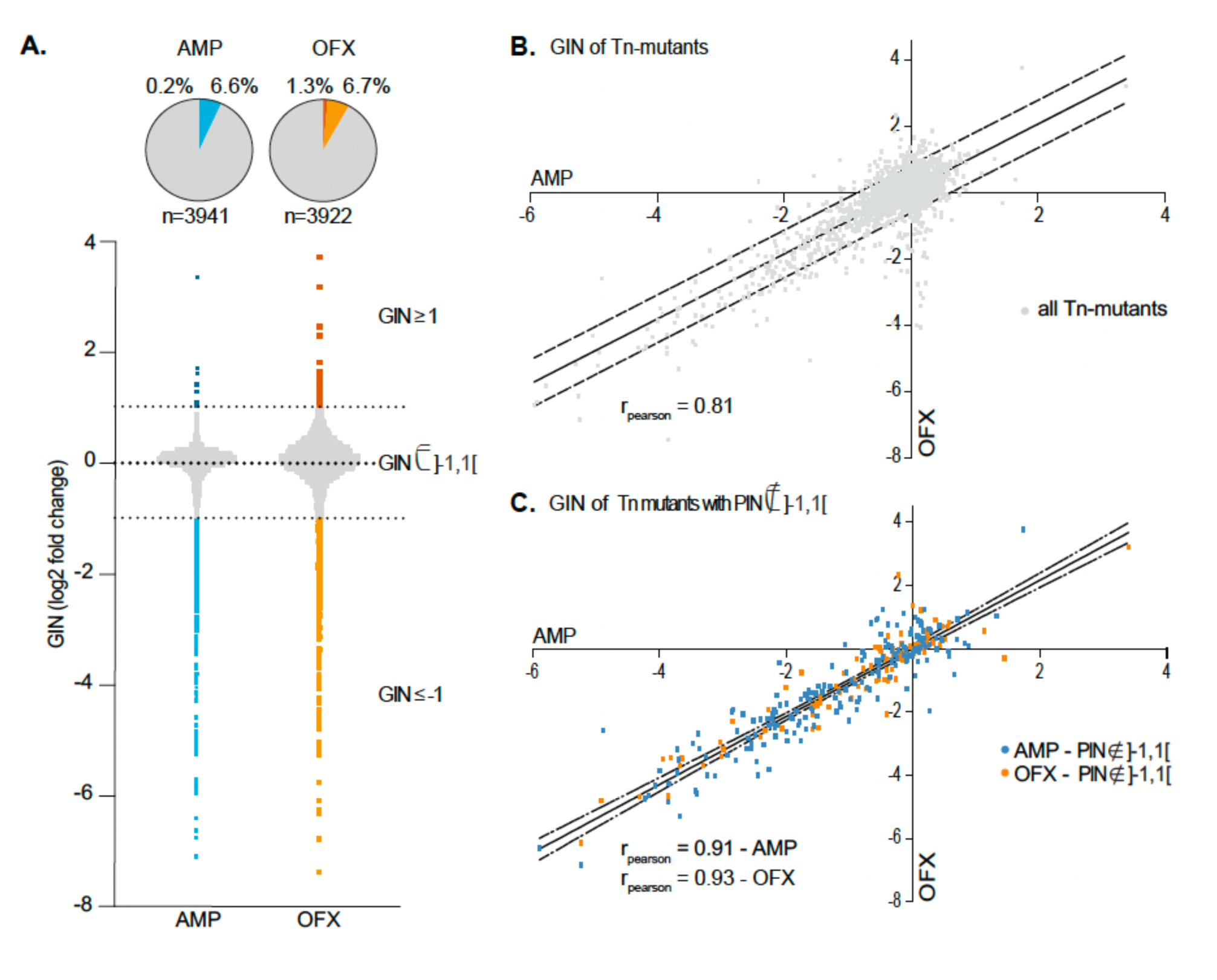
Genome-wide Tn-seq analysis of genes aIecting growth fitness. (**A**) GIN values for each non-essential gene under OFX and AMP conditions. Data are log_2_ fold change, significant thresholds were set at +1 and -1. (**B**) GIN values comparison of all non-essential Tn-mutants across the 2 conditions (AMP vs OFX). (**C**) GIN values comparison of all Tn-mutants with altered persistence (PIN ∉]-1, 1[) across experiments (AMP vs OFX). Curves represent the fitted linear regression with the 95% confidence intervals. (Blue = AMP; orange = OFX).

**Extended data Figure 3.**
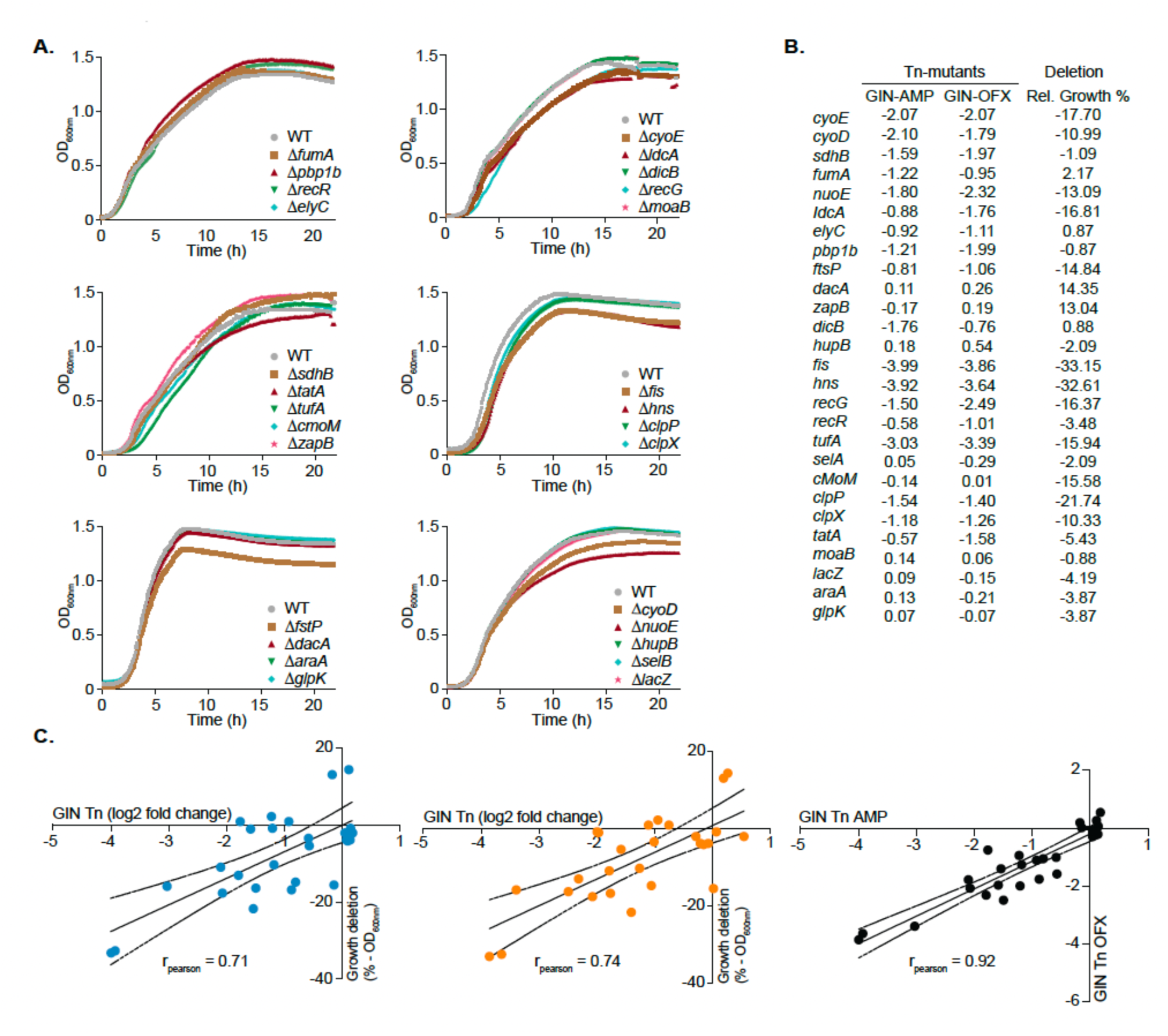
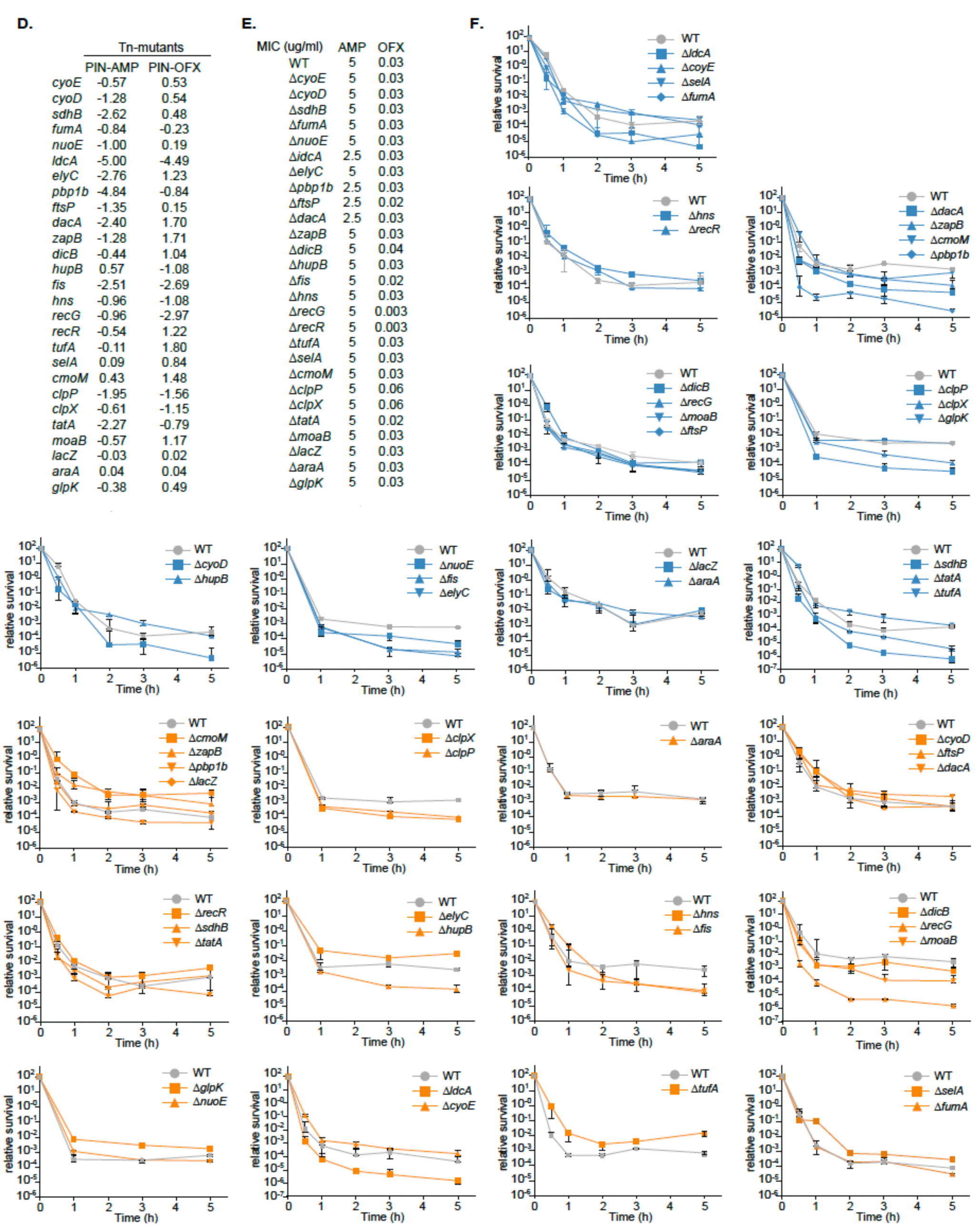
Characterization of the gene deletion mutants. (**A-C**) Growth characterization of the deletion strains. (**A**) Growth curve (OD_600nm_). (**B**) GIN values of Tn-mutants for the AMP and OFX conditions and relative growth of the corresponding deletion mutants, measured as the time required to reach an OD₆₀₀ of 0.8 and expressed relative to the WT strain (set at 100%). (**C**) Comparison of GIN values between the Tn-mutants and the relative growth of the corresponding deletion mutants for the AMP (blue) and OFX (orange) conditions, and comparison of GIN values between the 2 conditions. Curve represents the fitted linear regression with the 95% confidence intervals. (**D-F**) Persistence characterization of the deletion strains. (**D**). MIC (μg/mL) of AMP and OFX for each deletion mutants. (**E**) PIN values of the Tn-mutants corresponding to the deletion mutants under AMP and OFX conditions. (**F**) Time-kill curves of each deletion mutants treated with AMP and OFX for 5 h. (Blue = AMP, orange = OFX, grey = WT strain).

**Extended data Figure 4.**
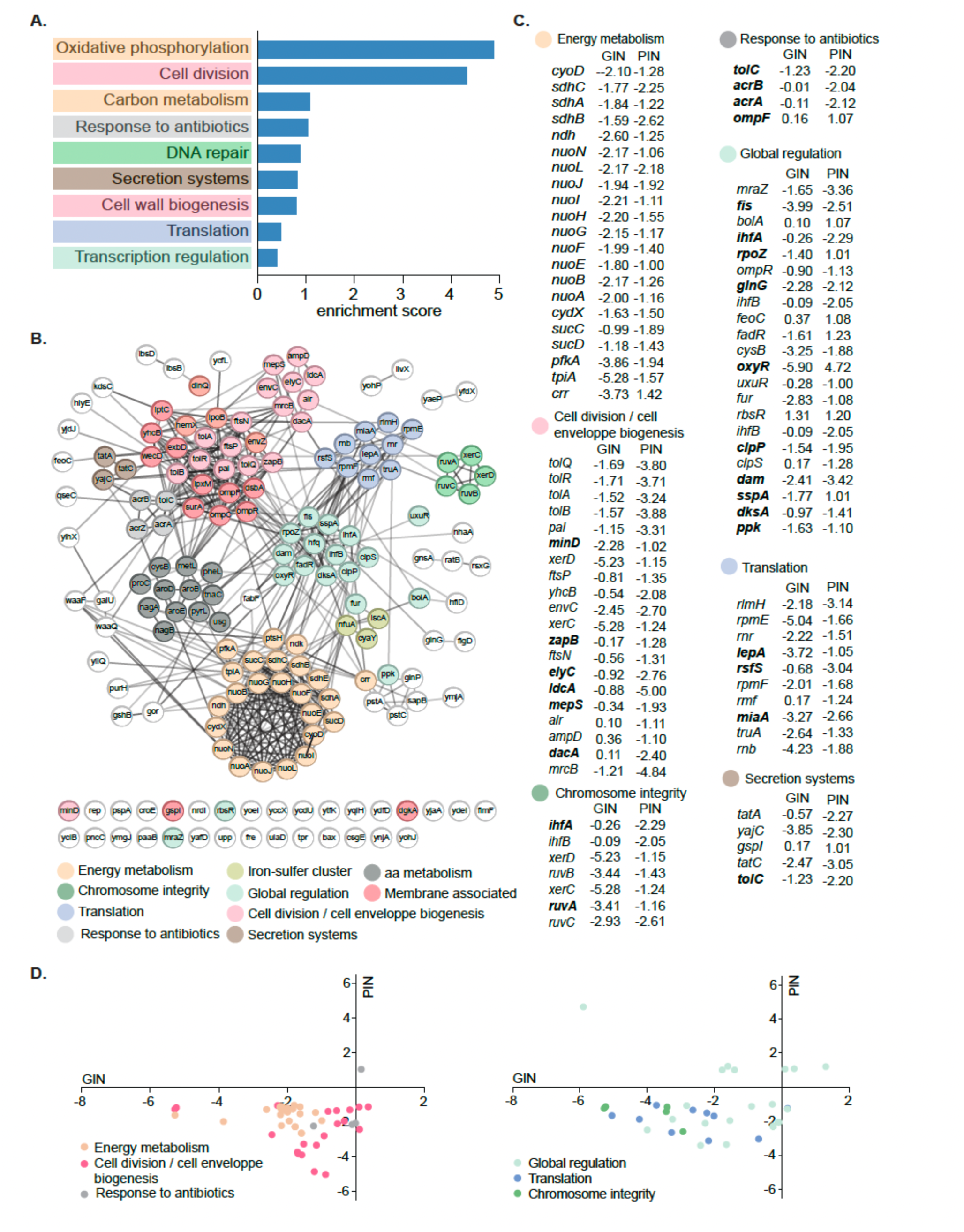
Functional network associated with persistence to ampicillin. (**A**) DAVID functional classification of genes involved in persistence to AMP. (**B**) STRING network analysis of genes associated with persistence to AMP. Only protein-coding genes are shown. Line thickness indicates the confidence of the association between proteins. Genes are organized and colored into functional modules based on their DAVID functional annotations and the connectivity between related pathways identified by STRING. (**C**) GIN and PIN values of genes in the main enriched functional modules. (**D**) Comparison of GIN and PIN values for genes within each functional module enriched under the AMP condition. Data represent log₂ fold changes derived from the Tn-seq analysis with colors indicating the corresponding functional modules.

**Extended data Figure 5.**
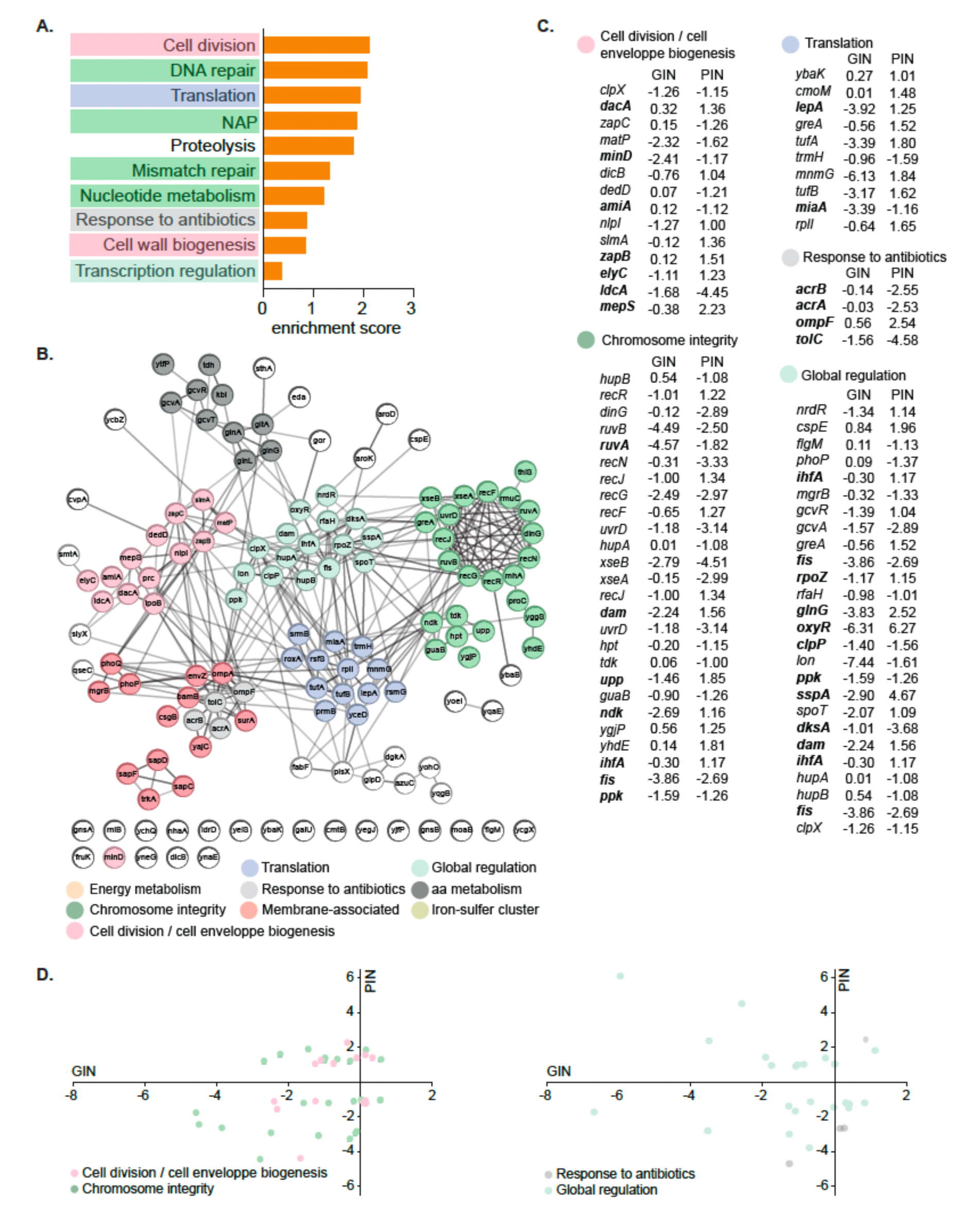
Functional network associated with persistence to ofloxacin. (**A**) DAVID functional classification of genes involved in persistence to OFX. (**B**) STRING network analysis of genes associated with persistence to OFX. Only protein-coding genes are shown. Line thickness indicates the confidence of the association between proteins. Genes are organized and colored into functional modules based on their DAVID functional annotations and the connectivity between related pathways identified by STRING. (**C**) GIN and PIN values of genes in the main enriched functional modules. (**D**) Comparison of GIN and PIN values for genes within each functional module enriched under the OFX condition. Data represent log₂ fold changes derived from the Tn-seq analysis with colors indicating the corresponding functional modules.

**Extended data Figure 6.**
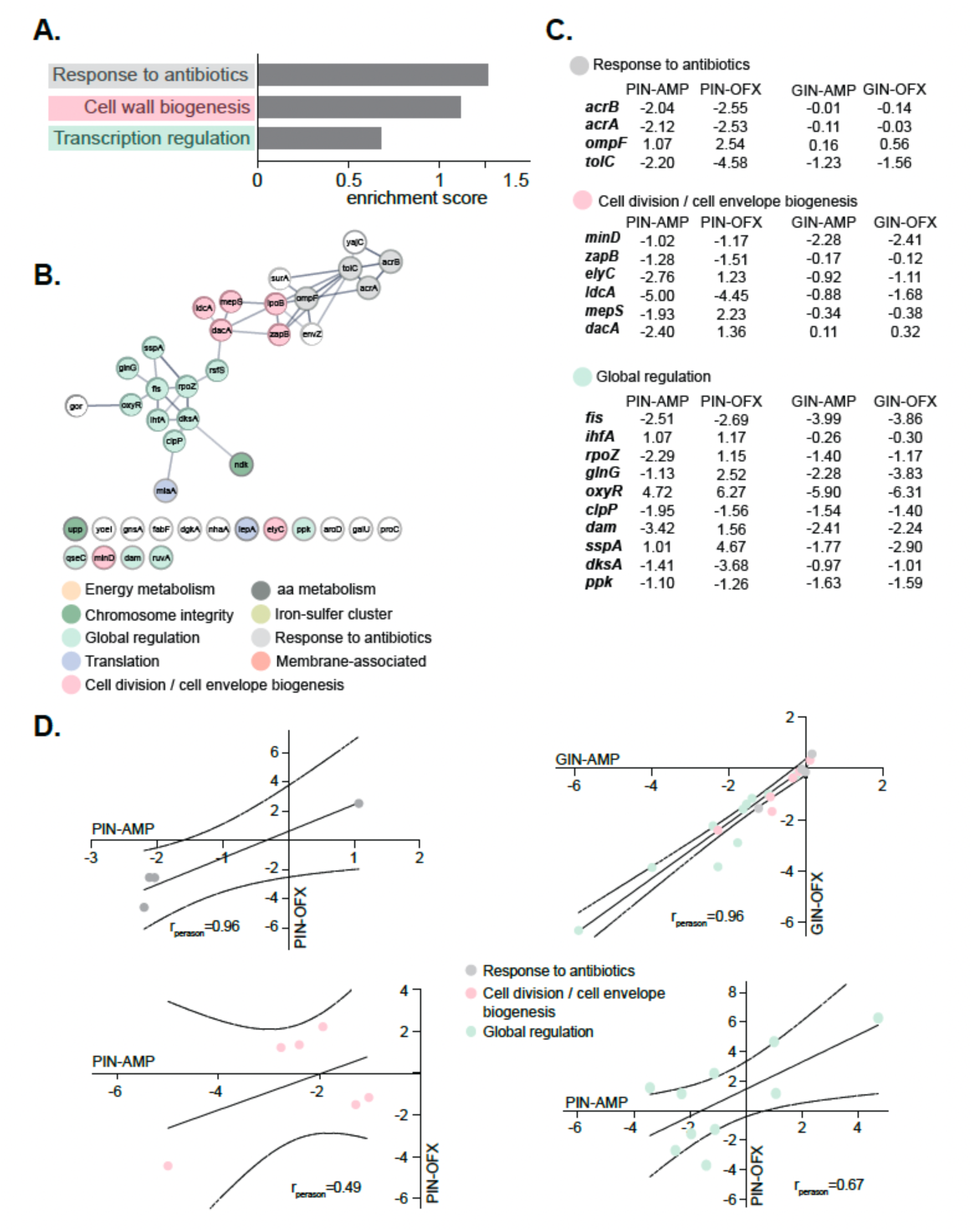
Functional network associated with persistence to both antibiotics. (**A**) DAVID functional classification of genes involved in persistence to both AMP and OFX. (**B**) STRING network analysis of genes associated with persistence to both antibiotics. Only protein-coding genes are shown. Line thickness indicates the confidence of the association between proteins. Genes are organized and colored into functional modules based on their DAVID functional annotations and the connectivity between related pathways identified by STRING. (**C**) GIN and PIN values of genes in the main enriched functional modules. (**D**) Comparison of PIN values for genes within each functional module between the two antibiotic conditions, and comparison of GIN values across all functional modules between the two conditions. Data represent log₂ fold changes derived from the Tn-seq analysis with colors indicating the corresponding functional modules.

